# Oxidative stress in alpha and beta cells as a selection criterion for biocompatible biomaterials

**DOI:** 10.1101/728683

**Authors:** Mireille M.J.P.E. Sthijns, Marlon J. Jetten, Sami G. Mohammed, Sandra M.H. Claessen, Rick de Vries, Adam Stell, Denise de Bont, Marten A. Engelse, Didem Mumcuoglu, Clemens A. van Blitterswijk, Patricia Y.W. Dankers, Eelco J.P. de Koning, Aart A. van Apeldoorn, Vanessa L.S. LaPointe

**Affiliations:** Department of Instructive Biomaterials Engineering, MERLN Institute for Technology-Inspired Regenerative Medicine, Maastricht University, Universiteitssingel 40, 6229 ER Maastricht, the Netherlands; Department of Complex Tissue Regeneration, MERLN Institute for Technology-Inspired Regenerative Medicine, Maastricht University, Universiteitssingel 40, 6229 ER Maastricht, the Netherlands; Department of Nephrology, Leiden University Medical Center, Albinusdreef 2, 2333 ZA Leiden, the Netherlands; Eindhoven University of Technology, P.O. Box 513, 5600 MB Eindhoven, the Netherlands; Hubrecht Institute, Uppsalalaan 8, 3584 CT Utrecht, the Netherlands

## Abstract

The clinical success of islet transplantation is limited by factors including acute ischemia, stress upon transplantation, and delayed vascularization. Islets experience high levels of oxidative stress due to delayed vascularization after transplantation and this can be further aggravated by their encapsulation and undesirable cell-biomaterial interactions. To identify biomaterials that would not further increase oxidative stress levels and that are also suitable for manufacturing a beta cell encapsulation device, we studied five clinically approved polymers for their effect on oxidative stress and islet (alpha and beta cell) function. We found that 300 poly(ethylene oxide terephthalate) 55/poly(butylene terephthalate) 45 (PEOT/PBT300) was more resistant to breakage and more elastic than other biomaterials, which is important for its immunoprotective function. In addition, PEOT/PBT300 did not induce oxidative stress or reduce viability in MIN6 beta cells, and even promoted protective endogenous antioxidant expression over 7 days. Importantly, PEOT/PBT300 is one of the biomaterials we studied that did not interfere with insulin secretion in human islets. These data indicate that PEOT/PBT300 may be a suitable biomaterial for an islet encapsulation device.

## Introduction

More than 40 million people worldwide suffer from type 1 diabetes (T1D), an autoimmune disease in which the pancreatic beta cells are destroyed, resulting in uncontrollable abnormal glycemic levels [1]. People with T1D need regular daily insulin injections and glucose monitoring to regulate their blood glucose, and they face a number of serious long-term secondary complications such as neuro- and retinopathy, kidney damage and cardiovascular disease. Severely affected patients with T1D are currently treated by a whole pancreas or clinical islet transplantation, but both interventions have their limitations, including limited donor availability, risks of unwanted comorbidities, the use of immunosuppressants to avoid rejection, and, in case of clinical islet transplantation (CIT), poor survival of the islets in the hepatic vasculature. At this moment, fewer than 1% of people with T1D undergo CIT, and in 60–70% of those who do, the transplanted islets eventually cease to function, and symptoms and complications associated with T1D return [2].

To increase the success of CIT, various biomaterial-based strategies are being considered. Open macroporous encapsulation devices that allow revascularization of islets have demonstrated some promise but it still takes at least 7–14 days until a new functional vasculature is established in transplanted islets [3] and immunosuppressants are always required. Macro- and microencapsulation of islets and beta cells are an alternative to open devices that introduce a physical barrier to protect the transplanted islets from the host’s immune system, potentially circumventing the need for immunosuppressive therapy [4] and mitigating some risks by protecting the patient from rogue cells in case of induced pluripotent- or embryonic stem cell-derived beta cell therapy [5-11]. Whatever the strategy chosen, the clinical success of encapsulation devices is hampered by a number of complex factors such as acute and long-term ischemia, limited vascularization and mass transport of crucial nutrients such as oxygen and insulin, and suboptimal biomaterial properties [12].

Indeed, the biomaterial used for manufacturing an islet encapsulation device requires careful selection [13]. The device itself should be suitable for handling during surgery, which means it should be compliant and resistant to breakage. Furthermore, scaffold stiffness has been shown to influence cell behavior by modulating the extracellular matrix and affecting the islet niche [14-16]. In addition, the biomaterial should be hydrophilic to facilitate insulin and glucose diffusion [17]. The general consensus is that in order to provide long-term support and protection of the islets from the patient and, conversely, to protect the patient from any dysfunctional cells, retrievable and non-degradable biomaterials are preferred. Apart from the aforementioned criteria, the manufacturing method and device design also dictate the selection of a biomaterial used for an islet encapsulation device. We have previously shown that a microwell scaffold platform comprising very thin porous polymer films chosen for their non-degradable, thermoplastic and mechanical properties separated individual islets from each other and supported their function and vascularization [18, 19].

One less-studied, but important factor to consider is the stress that biomaterials can impart on encapsulated islets and beta cells [20-22]. When in direct contact with cells, biomaterials can induce oxidative stress, which is known to decrease islet survival, and can diminish the success rate of CIT in the case of biomaterial-based beta cell replacement therapy [23-26]. During CIT, islets experience unusually high levels of oxidative stress in the first two weeks due their dissociation from the vasculature and deprivation of oxygen [27], a phenomenon that may be less prominent in open devices, but is a major issue in immunoprotective closed devices. It is also important to note that during the first onset of diabetes, oxidative stress can reduce the survival of the autoimmune-rejected islets [28, 29]. Oxidative stress occurs when the reactive oxygen species exceed (endogenous) antioxidants and the balance cannot be restored [20]. Beta cells are particularly sensitive to oxidative stress since they contain very low antioxidant levels, but little is known about the sensitivity of alpha cells [30]. Previously it has been shown that biomaterials can induce oxidative stress, which limits their biocompatibility [31].

In this study, we investigate the cell–biomaterial interaction using a series of different polymers with particular attention given to oxidative stress and islet function (determined by gene expression-related changes) in rodent pancreatic endocrine cells. All biomaterials studied are considered to be biocompatible based on past performance in *in vivo* studies and, in some cases, their current clinical use (Table 1). We hypothesize that different polymeric biomaterials can induce different levels of oxidative stress in the blood glucose–controlling pancreatic endocrine cells. We postulate that proper selection of a “beta cell–compatible biomaterial” used in beta cell replacement therapy should be based on a careful balance between basic biomaterials properties—allowing device fabrication, surgical handling, long-term structural support, and implant retrieval—and biomaterial-endocrine cell interactions leading to minimal, or no cell stress, providing especially the beta cells with the best head start possible to ensure long-term survival and function after transplantation.

**Table 1:** Biomaterials properties and applications.

| Biomaterial name | Code | Form* | Pore size (μm) | Thickness (μm) | Medical applications |
| --- | --- | --- | --- | --- | --- |
| Tissue culture polystyrene | TC | N/A | N/A | N/A | N/A |
| 300 polyethylene oxide terephthalate 55 polybutylene terephthalate 45 | PEOT/PBT300 | P | N/A | 20 | Bone filling, neural tissue engineering [32, 33] |
| Polyethylene terephthalate | PET | F | 0.2 | 8 | Vascular grafts, islet transplantation [34, 35] |
| Polyvinylidene fluoride | PVDF | F | 0.2 | 55 | Suture material, hernia meshes, bone and muscle tissue engineering [36, 37] |
| Ureidopyrimidinone-polycarbonate | UPy-PC | F | N/A | 80 | Pelvic floor repair [38] |
| 4000 polyethylene oxide terephthalate 30 polybutylene terephthalate 70 | PEOT/PBT4000 | P | N/A | 20 | Islet transplantation [18] |

\*F = film; P = pellets

Here we evaluated the physical–mechanical properties of five selected polymers, including hydrophobicity and elasticity, as well as whether they affected the viability of and supported angiogenesis in alpha (αTC1) and beta (MIN6) cells. We then studied the effect of the biomaterials on the intracellular level of oxidative stress and Nrf2-mediated endogenous antioxidant gene expression, and beta cell function–related gene expression. We found that some biomaterials induced significant oxidative stress, whereas others promoted the production of protective antioxidants. We observed that αTC1 and MIN6 cells responded differently to the polymers, and this response changed over time, which is important data to consider for a beta cell encapsulation device. Finally, islet function was determined by a glucose-stimulated insulin secretion test. With these criteria, we identified PEOT/PBT300 as having suitable properties for use in an islet encapsulation device.

## Material and methods

### Polymer films

For the five polymers, 8–80 µm-thick films were used in this study. PET was purchased from GVS (Lancester, United Kingdom) and PVDF from Thermo Fisher Scientific (Waltham, United States). PEOT/PBT4000 and PEOT/PBT300 (Polyvation BV, Groningen, the Netherlands) were prepared as previously published [18] by film casting on an automated film applicator (Elcometer 4340, Elcometer BV, Utrecht, the Netherlands). In short, 15% (w/w) PEOT/PBT solutions were prepared in a mixture of chloroform and 1,1,1,3,3,3-hexafluoro-2-isopropanol, at respective ratios of 65:35 (w/w) for PEOT/PBT4000 and 90:10 (w/w) for PEOT/PBT300. Subsequently, the polymer solutions were casted at ambient temperature and humidity (10%) with an automated film applicator at a casting speed of 5 mm/s and initial thickness of 250 μm. After casting, the polymer films were dried overnight under a nitrogen stream followed by overnight incubation in ethanol to remove all residual solvents. Finally, the films were air dried. Ureidopyrimidinone-polycarbonate UPy-PC films were prepared by drop-casting into Teflon molds. Chain-extended ureidopyrimidinone (UPy)-based polycarbonate (CE-UPy-PC, SupraPolix BV, Eindhoven, the Netherlands) was dissolved in chloroform/hexafluoro-2-isopropanol (95/5%) to a final concentration of 20 mg/mL and casted on a 4 × 10.5 cm Teflon mold and dried overnight. Films were subsequently removed from the mold, transferred to a petri dish and dried overnight in a vacuum oven at ambient temperature. Tissue culture polystyrene used as a reference material was purchased from Thermo Fisher Scientific (Waltham, United States).

### Biomaterials properties

The mechanical properties of each biomaterial were determined using an Electroforce (3230-ES Series III) equipped with a 45/450 kN load cell according to ASTM Standard D882-02. The dimensions of each biomaterial film were 35 × 10 mm, and the effective area between the clamps was 15 × 10 mm, except for PEOT/PBT300 and PEOT/PBT4000, which had an effective area of 10 × 10 mm and 5 × 10 mm, respectively, due to the travel limits of the machine. The ramp rate was set at a strain of 1%/min. Each biomaterial was tested three times both parallel and perpendicular to the film casting direction. Stress-strain curves (including peak stress, failure stress, and failure strain) were measured, and the Young’s Modulus was determined by calculating the slope within the proportionality limit of the curve.

### Water contact angle

The static sessile drop method was used to measure water contact angles for each of the biomaterials to determine their hydrophobicity. A drop shape analyser (Kruss, DSA25S) and Drop Shape Analysis 4 software was used to perform 14 separate measurements per biomaterial type.

### Scanning electron microscopy (SEM) imaging

Scanning electron microscopy (SEM) imaging was used to evaluate cell morphology on the biomaterials. The biomaterials alone and MIN6 cells grown for 1 or 7 days on these biomaterials were evaluated. Cells were fixed using 3.6% (v/v) formalin in PBS for 30 min at ambient temperature. All samples were fixed on stubs using carbon tape and sputter coated with gold for 60 s using a Cressington sputter coater. Samples were evaluated using a FEI Teneo microscope under high vacuum and secondary electron mode. Images of the biomaterials were taken at 1000 and 15000 times magnification, while images of the cells on the biomaterials were taken at a 500, 2500 and 15000 times magnification.

### Cell culture

Mouse alpha cells (αTC1 Clone 6; ATCC CRL-2934), passage 12–19, were cultured in DMEM (Sigma-Aldrich D6046) supplemented with 10% (v/v) fetal bovine serum, 15 mM HEPES, 0.1 mM non-essential amino acids, 1.5 g/L sodium bicarbonate, and 2.0 g/L glucose. Mouse beta cells (MIN6) were kindly provided by Dr. Caroline Arous from the Wehrle-Haller laboratory (University of Geneva, Switzerland). MIN6 clone b1 cells, passage 34–42, were cultured in DMEM (Sigma-Aldrich D6046) supplemented with 10% (v/v) fetal bovine serum, 10 mM HEPES, 1 mM sodium pyruvate, 2.0 g/L glucose, and 50 μM 2-mercaptoethanol. All cells were cultured in a humidified atmosphere containing 5% CO_2_ at 37°C.

### Human islet retrieval

Human islets were obtained from Prodo Laboratories Inc. (Aliso Viejo, USA) and Leiden University Medical Center (LUMC, Leiden, the Netherlands) from 5–6 different donors (two male, four female). Pancreatic islet isolation at LUMC was performed as previously described [13]. Human donor islets were used if deemed unsuitable for CIT and if research consent was present according to national laws and regulations. The average age, body mass index and islet equivalent quantity (IEQ) of the donors were 46 ± 15 years, 26.5 ± 3.87 and 20286 ± 22170, respectively. Islets from people with T1D (defined as an HbA1c > 7.0%) and a trauma that included pancreatic damage were excluded from analysis.

### Cell seeding

In preparation for cell seeding, biomaterials were punched into circular samples with a diameter of approximately 0.7 cm or 1.55 cm and washed in ethanol overnight. Vaseline was used to adhere the biomaterials into the wells of a 96-well (for oxidative stress experiments), or 24-well (for gene expression) plate. Cells were seeded at a density of 4.4 × 10^5^ cells/cm^2^ and were cultured for 1 or 7 days.

### Oxidative stress assay

To measure intracellular oxidative stress, culture medium was removed, and the cells were preincubated at 37°C in 5% CO_2_ for 45 min in 20 μM DCFH-diacetate in exposure medium comprising minimal essential Dulbecco’s modified eagle medium (Gibco 11880-028) supplemented with 15 mM HEPES, 0.1 mM non-essential amino acids, 1.5 g/L sodium bicarbonate, 3.0 g/L glucose and 4 mM L-glutamine. A 30-min incubation in 200 or 400 μM (for αTC1 and MIN6 cells, respectively) hydrogen peroxide (H_2_O_2_) was used as a positive control to induce oxidative stress. Cells were washed in phosphate-buffered saline (PBS), and the probe fluorescence was measured over a period of 1 h on a CLARIOstar microplate reader (BMG Labtech, Cary, North Carolina, United States) with an excitation wavelength of 485 nm and emission at 538 nm. The area under the curve was considered the total oxidative stress experienced by cells and was normalized to the number of viable cells and a reference sample of cells on tissue culture polystyrene after 1 day.

### Viability

To normalize oxidative stress to cell number, viability was determined with a CellTiter-Glo 2.0 Cell Viability Assay (Promega, Madison, United States) according to the manufacturer’s instructions. After 10 min of incubation, luminescence was measured for a couple of minutes on a CLARIOstar microplate reader. Viability was calculated as percentage relative to controls.

### Gene expression

H_2_O_2_ was removed from cell cultures after 30 min and replaced with fresh culture medium. After 2.5 h, total RNA was isolated using the RNeasy Micro Kit (Qiagen, Hilden, Germany). The RNA quantity and quality were assessed on a BioDrop μLITE+ (BioDrop, Cambridge, United Kingdom). Complementary DNA (cDNA) was made by converting at least 100 ng RNA with the iScript cDNA synthesis kit (Bio-Rad). Quantitative PCR (qPCR) was done in a 20 μl reaction using the iQ SYBR Green Supermix on a Real-Time PCR Detection System (Bio-Rad). Samples were incubated for 3 min at 95°C and the thermocycling was 12 s at 95°C and 30 s at 58°C for 38 cycles. Validated primers were used at a concentration of 300 nM (Table 2). Hypoxanthine guanine phosphoribosyl transferase (*Hprt*) was used as a housekeeping gene. Relative gene expression was determined using the Livak (2^− ΔΔCT^).

**Table 2:**
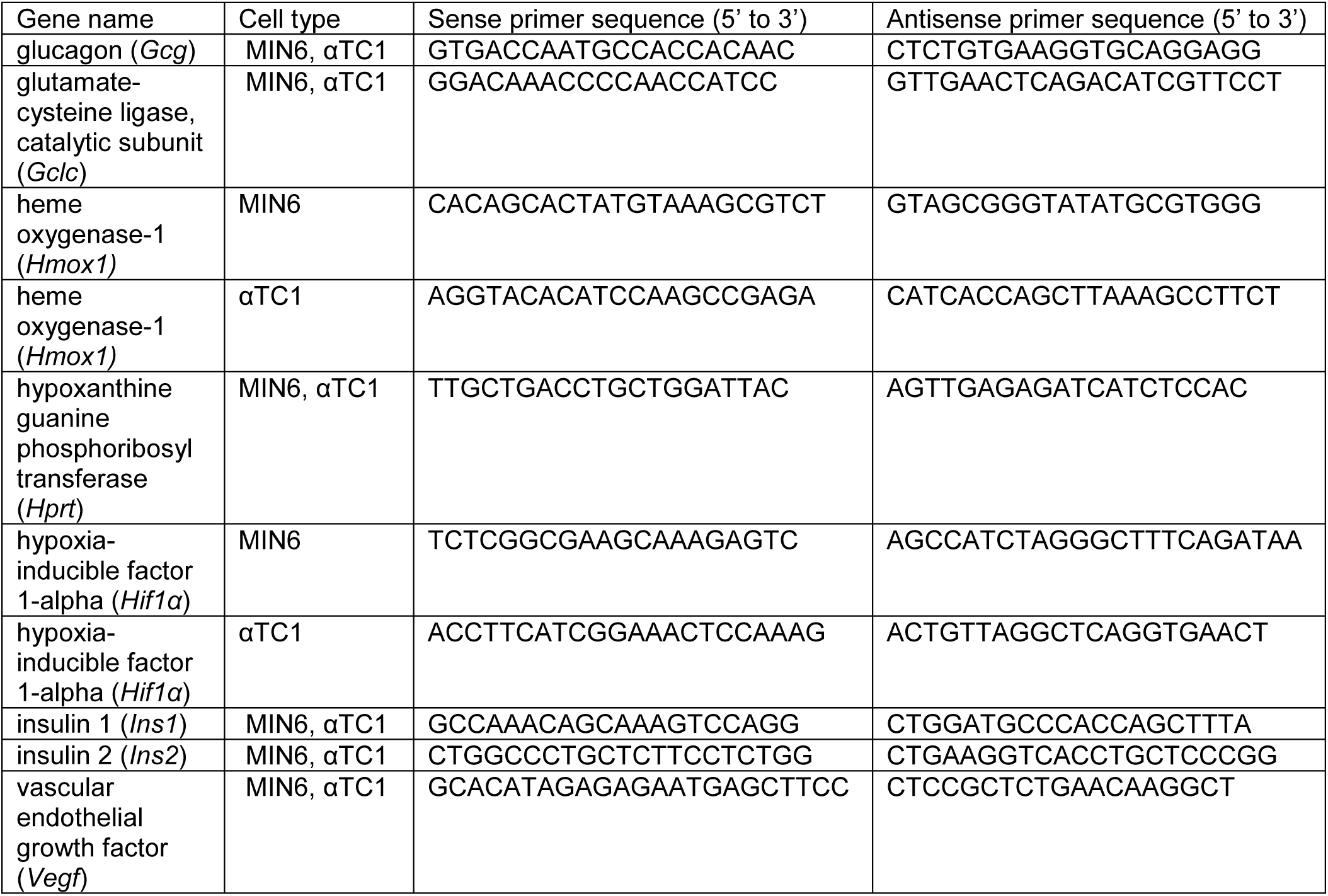
Primer sequences.

### Glucose-stimulated insulin secretion

To prepare donor islets for the experiments, the transport medium was carefully removed immediately upon their arrival and replaced with CMRL-1066 medium (Pan-Biotech, Aidenbach, Germany) supplemented with 5.5 mM glucose, 26 mM NaHCO_3_, 2 mM GlutaMax, 50 μg/ml penicillin-streptomycin, 10 μg/ml ciprofloxacine and 10% fetal bovine serum. The IEQ number for each donor was determined with dithizone staining (Lonza, Basel, Switzerland), and 50 IEQ in 1 ml was used for each sample [18]. The biomaterials were placed in 12 μm pore size cell culture inserts (Millipore), after which islets were seeded on top of each biomaterial and cultured for 7 days. Islets in a non-adhesive, 24-well plate (Greiner Bio-one, Vilvoorde, Belgium) were used as controls. To measure the glucose-stimulated insulin secretion, islets were sequentially exposed to 1.67 mM (low), 16.7 mM (high) and 1.67 mM (low) glucose in filtered Krebs buffer stock solution, pH 7.3–7.5, for 1 h each. After each incubation step, the supernatant was removed, and the sample was centrifuged at 300 × *g* for 3 min at ambient temperature. The supernatant was transferred to a microcentrifuge tube and stored at −20°C. Insulin secretion was determined with a human insulin ELISA kit (Mercodia, Uppsala, Sweden) according to the manufacturer’s instructions. After the final incubation step, the cells were lysed and the amount of DNA was determined with a Quant-iT PicoGreen dsDNA Assay (Thermo Fisher Scientific, Waltham, United States). The total amount of insulin released in each incubation step per sample was displayed relative to the DNA quantity, and the glucose stimulation indices were calculated by dividing the insulin secretion in high glucose medium by the basal insulin secretion in low glucose medium.

### Statistics

All data are presented as mean ± SEM from at least 2 technical duplicates and three independent experiments (n ≥ 3). Independent samples with equal variances were assessed for statistical significance with a *t*-test. P values < 0.05 were considered statistically significant.

## Results

### PEOT/PBT block copolymers have desirable physical elastic and hydrophilic properties

Several thermoplastic polymers were preselected on the basis of their past performance and, in some cases, their clinical use (Table 1). The ideal biomaterial for an encapsulation device would be elastic for implantation and hydrophilic to enhance insulin and glucose diffusion [17]. Thus, we characterized the mechanical strength, elasticity, and hydrophilicity of the five polymers. Tensile testing was performed with the biomaterials in the directions parallel (Figure 1) and perpendicular (Supplementary Figure 1) to film casting. Both directions (Supplementary Table 1–2) showed similar statistically significant results. For peak stress, PET and PEOT/PBT4000 had the highest values of 14.4 and 14.7 MPa, respectively (Figure 1A). PET had a statistically significantly higher Young’s modulus (7.8 MPa) and failure stress (14.2 MPa) compared to the other four biomaterials (Figure 1B–C). In addition, the failure strain of PEOT/PBT4000 (429.0%) and PEOT/PBT300 (547.3%) was more than two times higher than the other biomaterials (92.9%), indicating that PEOT/PBT300 and PEOT/PBT4000 were elastic and resisted higher strains than PET (Figure 1D). Next, the water contact angle of the biomaterials revealed that PVDF was more hydrophobic (131.4°) compared to all other biomaterials (ranging from 61.5° to 82.0°; Figure 1E). The surface structure of the different biomaterials was different (Supplementary Figure 2) but the morphology of MIN6 cultured on the biomaterials did not change (Supplementary Figure 3) apart from fewer cells adhering to PEOT/PBT4000. Together, these findings indicate that PEOT/PBT300 and PEOT/PBT4000 are good candidates for the encapsulation device because of their elasticity and resistance to breakage, while PET should be excluded from consideration. PVDF is also a less favorable biomaterial due to its hydrophobicity.

**Figure 1:**
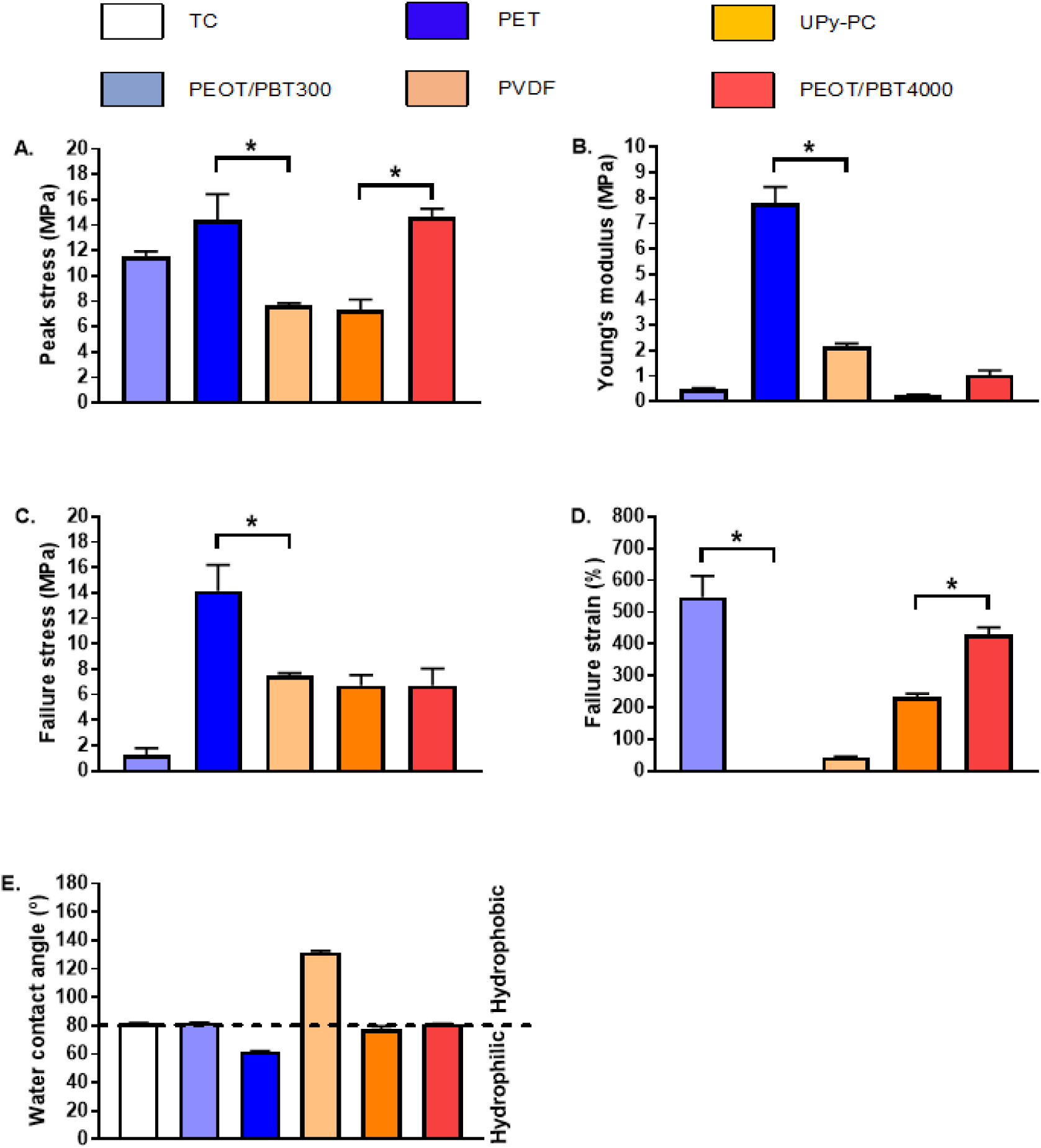
Physical–mechanical and hydrophobic properties of five selected polymers. Tensile testing in the direction parallel to casting revealed that PET and PEOT/PBT4000 had higher peak stress than the other biomaterials (A). The Young’s modulus and failure stress of PET was higher than the other biomaterials (B–C). PC and PEOT/PBT4000 had the highest failure strain (D). PVDF was the most hydrophobic and PET was most hydrophilic biomaterial (E). *N*=3 and data are presented as mean ± SEM; * *p* ≤ 0.05.

### Vegf expression was not affected by any biomaterials tested

In addition to the physical properties of the biomaterial, its ability to support angiogenesis is important for determining the best-suited biomaterial for an islet encapsulation device. It is known that the biomaterial composition is essential to induce angiogenesis [39, 40]. To verify whether the biomaterials induce angiogenesis pathways in MIN6 and αTC1 cells, *Vegf* and *Hif1α* expression were measured by qPCR and given as relative expression compared to *Hprt1*, an internal housekeeping gene (Figure 2). *Hif1α* is induced by low oxygen and is a transcription factor that mediates the transcription of *Vegf*, which enhances angiogenesis by increasing endothelial cell sprouting. *Hif1α* expression in MIN6 cells cultured on UPy-PC and PEOT/PBT4000 was significantly decreased at day 1, whereas at day 7 *Hif1α* expression in MIN6 cells was decreased when cultured on all five biomaterials, compared to that in cells cultured on the reference biomaterial, tissue culture polystyrene (TC; Figure 2A–B). For αTC1 cells, *Hif1α* expression was significantly decreased on PET at day 1 and day 7 compared to TC (Figure 2C– D). In comparison, *Vegf* expression was significantly decreased in MIN6 and αTC1 cells cultured on PEOT/PBT300 at day 1, MIN6 cells on PEOT/PBT4000 at day 1, compared to TC (Figure 2E–G). By day 7, however, *Vegf* expression was similar to control TC in both MIN6 and αTC1 cells (Figure 2F–H). These results indicate that, by day 7, all biomaterials induced a similar *Vegf* expression as the reference biomaterial TC, and that the induced *Vegf* expression was not mediated by *Hif1α* transcription.

**Figure 2:**
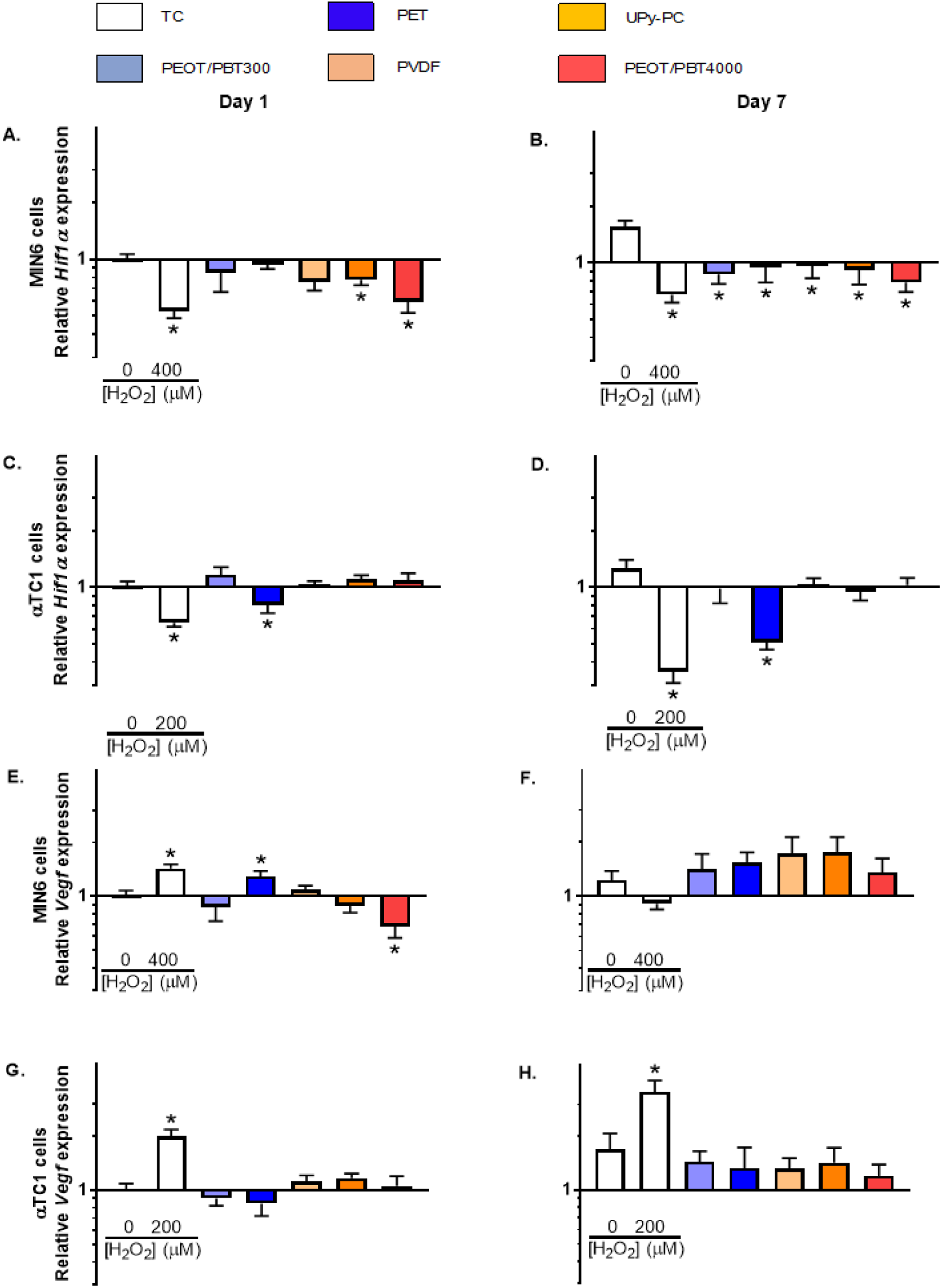
Gene expression levels of angiogenic-related proteins (*Hif1α* and *Vegf*) of MIN6 and αTC1 cells cultured on different polymer films. *Hif1α* expression was decreased in MIN6 cells cultured on UPy-PC and PEOT/PBT4000 on day 1 (A) and on all biomaterials on day 7 (B). *Hif1α* expression in αTC1 cells was decreased on day 1 when cultured on PET (C) and on day 7(D). *Vegf* transcript expression was significantly lower on day 1 in MIN6 cells cultured on PEOT/PBT4000 compared to TC (E), whereas there was no difference on day 7 (F). For αTC1 cells, the *Vegf* transcript expression was significantly lower in cells cultured on PEOT/PBT300 and PET on day 1 compared to TC (G), whereas there was no significant difference on day 7 (H). The positive control hydrogen peroxide (H_2_O_2_) decreased *Hif1α* expression in all conditions (A–D), while *Vegf* expression was increased (E,G–H), except in MIN6 cells at day 7 where *Vegf* expression was decreased (F). *N*=3 and data are presented as mean ± SEM relative to the housekeeping gene *Hprt* and the expression on TC at day 1; * *p* ≤ 0.05.

### Some biomaterials induce oxidative stress, which is cell type- and time-dependent

We sought a biomaterial that would not induce oxidative stress. We therefore measured intracellular oxidative stress by the DCFH assay in MIN6 and αTC1 cells cultured for 7 days on the different biomaterials. Cells cultured on TC in the absence and presence of H_2_O_2_ were used as negative and positive controls for oxidative stress, respectively. Overall, each different biomaterial induced a different level of oxidative stress. In addition, further differences in oxidative stress could be assigned to the different cell type or culture period. PEOT/PBT300 did not induce a detectable increase in oxidative stress at day 1 or 7 compared to the negative control in either cell type.

In MIN6 cells, only UPy-PC and PEOT/PBT4000 significantly increased the oxidative stress to 211% and 309% at day 1 compared to the negative control (Figure 3A). The increase in oxidative stress induced by PEOT/PBT4000 in MIN6 cells was similar to that induced by the positive control (TC with 400 μM H_2_O_2_) at day 1 (Figure 3A). By day 7, the elevated levels of oxidative stress in MIN6 cells on UPy-PC and PEOT/PBT4000 decreased compared to day 1 (Figure 3B). Interestingly, MIN6 cells on the positive control also had reduced oxidative stress at this time point, suggesting they had activated an endogenous protective mechanism.

**Figure 3:**
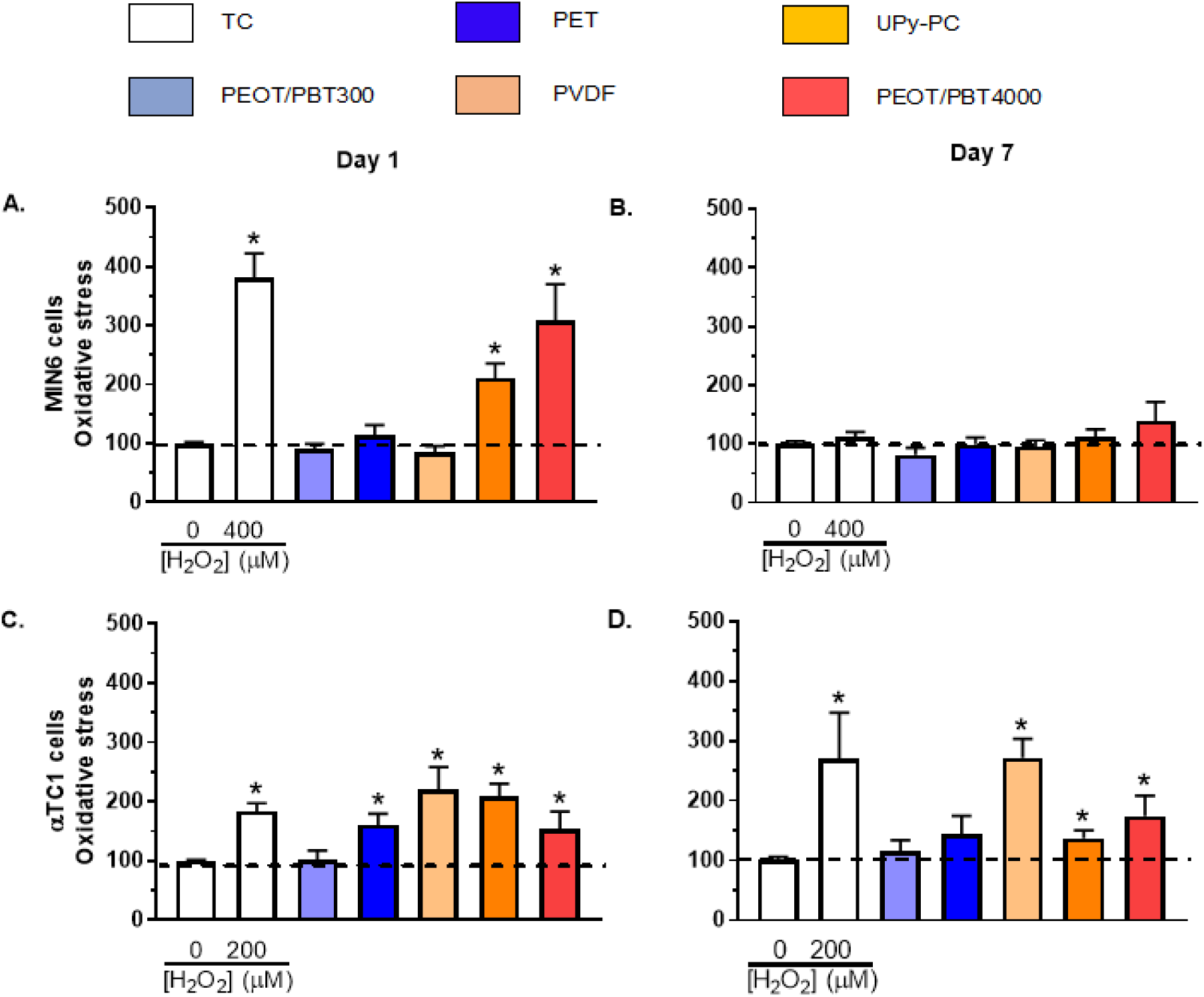
Oxidative stress levels of MIN6 or αTC1 cells cultured on different polymer films. MIN6 cells had a significantly increased level of oxidative stress on both UPy-PC and PEOT/PBT4000 on day 1 compared to TC (A), whereas neither hydrogen peroxide (H_2_O_2_) nor the biomaterials induced oxidative stress on day 7 (B). In αTC1 cells, 200 μM of H_2_O_2_ induced a significant increase in oxidative stress, as did all biomaterials except for PEOT/PBT300 (C). Unlike in MIN6 cells, no decrease in oxidative stress was measured in αTC1 cells exposed to H_2_O_2_ or cultured on biomaterials on day 7 (D). *N*=3 and data are presented as mean ± SEM compared to the oxidative stress measured in MIN6 or αTC1 cells on TC at day 1; * *p* ≤ 0.05.

In αTC1 cells, all biomaterials except for PEOT/PBT300 induced a significant increase in oxidative stress at day 1 compared to the negative control, with levels similar to the positive control (TC with 200 μM H_2_O_2_; Figure 3C). In contrast to what was observed in MIN6 cells, αTC1 cells at day 7 showed no significant reduction in the level of oxidative stress induced by the different biomaterials or the positive control when compared to day 1 (Figure 3D), suggesting they may lack the endogenous protective mechanism found in MIN6 cells. Notably, αTC1 showed sensitivity to oxidative stress induced by four of the five polymers over time (all except PEOT/PBT300), whereas MIN6 cells were sensitive to oxidative stress imposed by two biomaterials (UPy-PC and PEOT/PBT4000) but only for 1 day.

### MIN6 cells express endogenous antioxidants coinciding with diminished oxidative stress

Cells have their own protective mechanisms against oxidative stress, and so we measured the expression of important regulators within the endogenous antioxidant systems to see how they were affected by interaction with the polymers. While we sought to find a biomaterial that did not induce oxidative stress in itself, it could be even more advantageous to find one that protects cells from other sources of induction. Gene expression of three of these regulators, *Hmox1, Gclc*, and *Nfe2l2* (a transcription factor regulating *Hmox1* and *Gclc* expression) were measured with qPCR relative to *Hprt1*, and compared to cells cultured on TC after 1 day (Figure 4).

**Figure 4:**
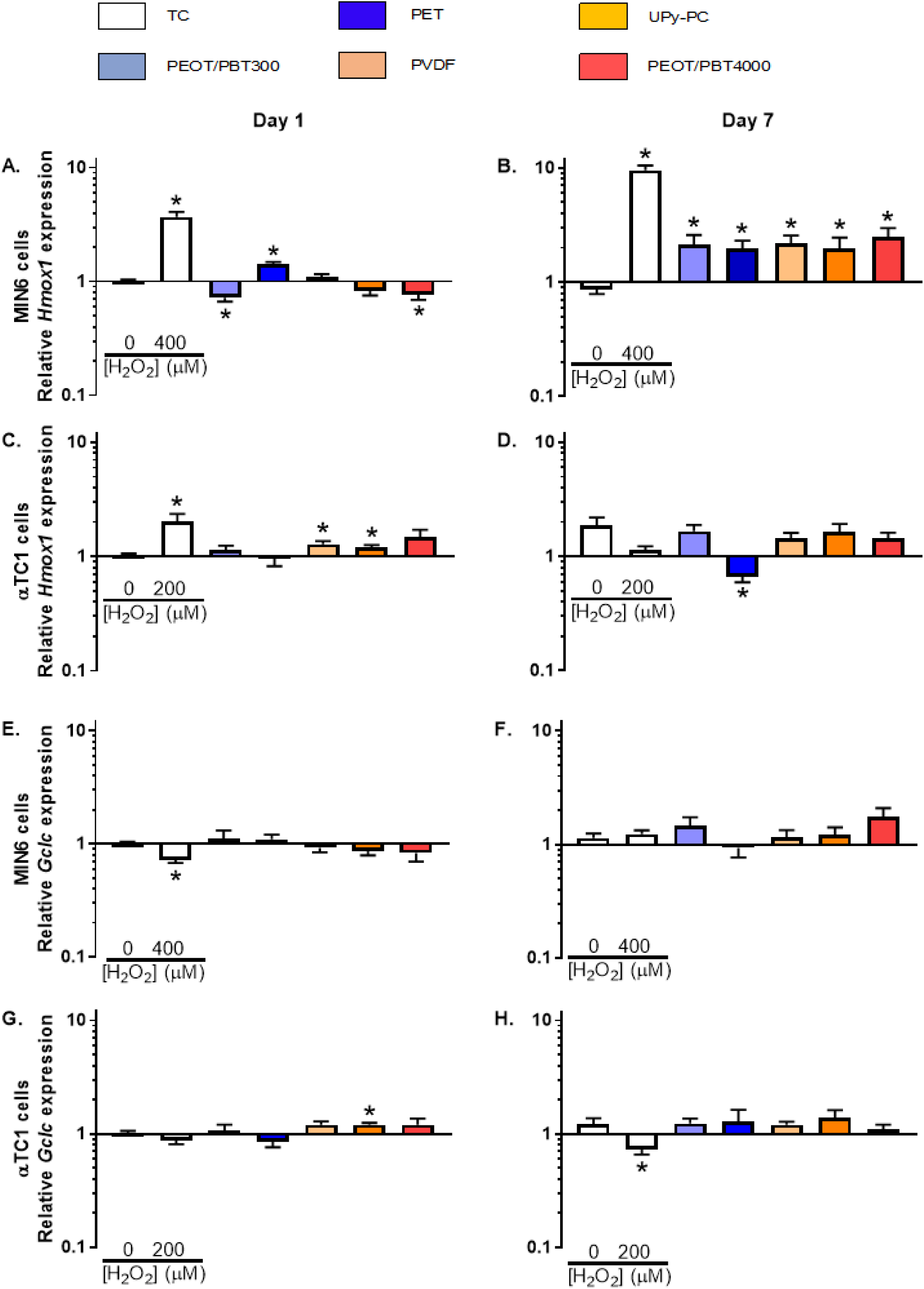
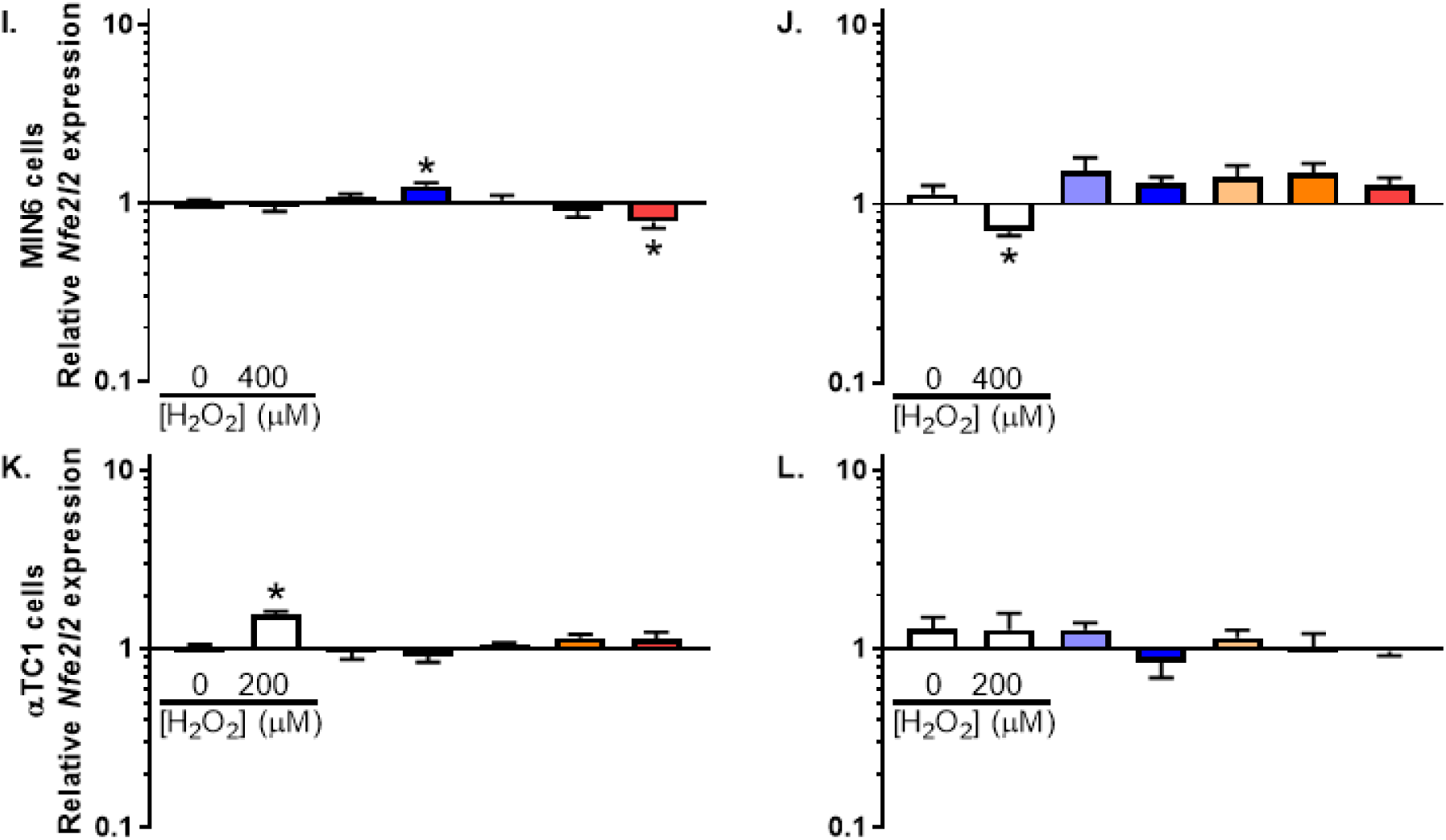
Gene expression levels of antioxidant proteins (*Hmox1, Gclc* and *Nfe2l2*) of MIN6 and αTC1 cells cultured on different polymer films. On day 1, *Hmox1* transcript expression was significantly lower in MIN6 cells cultured on PEOT/PBT300 and PEOT/PBT4000 compared to TC, while culturing on PET increased expression (A). On day 7, *Hmox1* expression was increased in MIN6 cells cultured on all biomaterials (B). In αTC1 cells on day 1 PVDF and UPy-PC increased *Hmox1* expression (C). On day 7, only PET decreased *Hmox1* expression, whereas culturing on the other biomaterials had no effect (D). In MIN6 cells, *Gclc* expression was not affected by all five biomaterials on day 1 (E) and day 7 (F). In αTC1 cells, UPy-PC increased *Gclc* expression on day 1 (G), whereas at day 7 no significant change was measured (H). *Nfe2l2* expression showed a similar effect as *Hmox1* expression (I-L). Hydrogen peroxide increased *Hmox1* expression, except for day 7 in αTC1 cells. Hydrogen peroxide decreased *Gclc* in MIN6 on day 1 and in αTC1 cells on day 7, while *Nfe2l2* expression was decreased in MIN6 cells due to exposure to hydrogen peroxide and increased in in αTC1 cells on day 1. *N*=3 and data are presented as mean ± SEM relative to the housekeeping gene *Hprt* and the expression on TC at day 1; * *p* ≤ 0.05.

On day 1, *Hmox1* expression in MIN6 was significantly lower on PEOT/PBT300 and PEOT/PBT4000 (0.72 and 0.77, respectively, compared to the control set to 1.00 on a log scale), compared to the negative control (Figure 4A). After 7 days, MIN6 cells cultured on all biomaterials showed significantly increased *Hmox1* expression, indicating the activation of the endogenous antioxidant system. The mean *Hmox1* expression of MIN6 cells cultured on the biomaterials was higher compared to that on TC (2.1 v. 0.86; Figure 4B). This increase in *Hmox1* gene expression was accompanied by a decrease in oxidative stress at day 7 compared to day 1 (Figure 3B). In αTC1 cells, only PET induced lower *Hmox1* expression at day 7 (Figure 4D).

*Gclc* expression was upregulated in MIN6 cells cultured on PVDF, UPy-PC, and PEOT/PBT4000 at day 7 compared to day 1, indicating an activation of antioxidant systems over time (Figure 4E–F). In αTC1 cells, *Gclc* expression was only increased at day 7 compared to day 1 when culturing αTC1 on PET (Figure 4G–H). *Nfe2l2* was only upregulated in MIN6 cultured on PET at day 1 (Figure 4I–L), indicating that *Nfe2l2* expression was not affected.

### Some biomaterials decreased viability

Oxidative stress is known to decrease cell viability [41], so we tested the effect of the cell-biomaterial interaction on the viability of MIN6 and αTC1 cells (Figure 5). The viability of cells cultured on TC with and without H_2_O_2_ (an inducer of oxidative stress) was used as reference samples. In MIN6 cells, UPy-PC and PEOT/PBT4000 significantly decreased viability by 32% and 82%, respectively, at day 1 compared to control TC (Figure 6A); MIN6 viability remained decreased (69% compared to control TC) on PEOT/PBT4000 at day 7 (Figure 6B). For αTC1 cells at day 1, UPy-PC, PEOT/PBT4000 and PEOT/PBT300 all decreased viability by 45%, 71% and 18%, respectively. At day 7, PVDF, UPy-PC and PEOT/PBT4000 decreased αTC1 cell viability by 43%, 32% and 63%, respectively (Figure 6C–D). We noted that the ability of MIN6 to increase *Hmox1* endogenous antioxidant protection over time was correlated to their viability on that specific biomaterial while the αTC1 cells that were unable to induce *Hmox1* endogenous antioxidant protection, did not show that correlation (Supplementary Figure 4).

**Figure 5:**
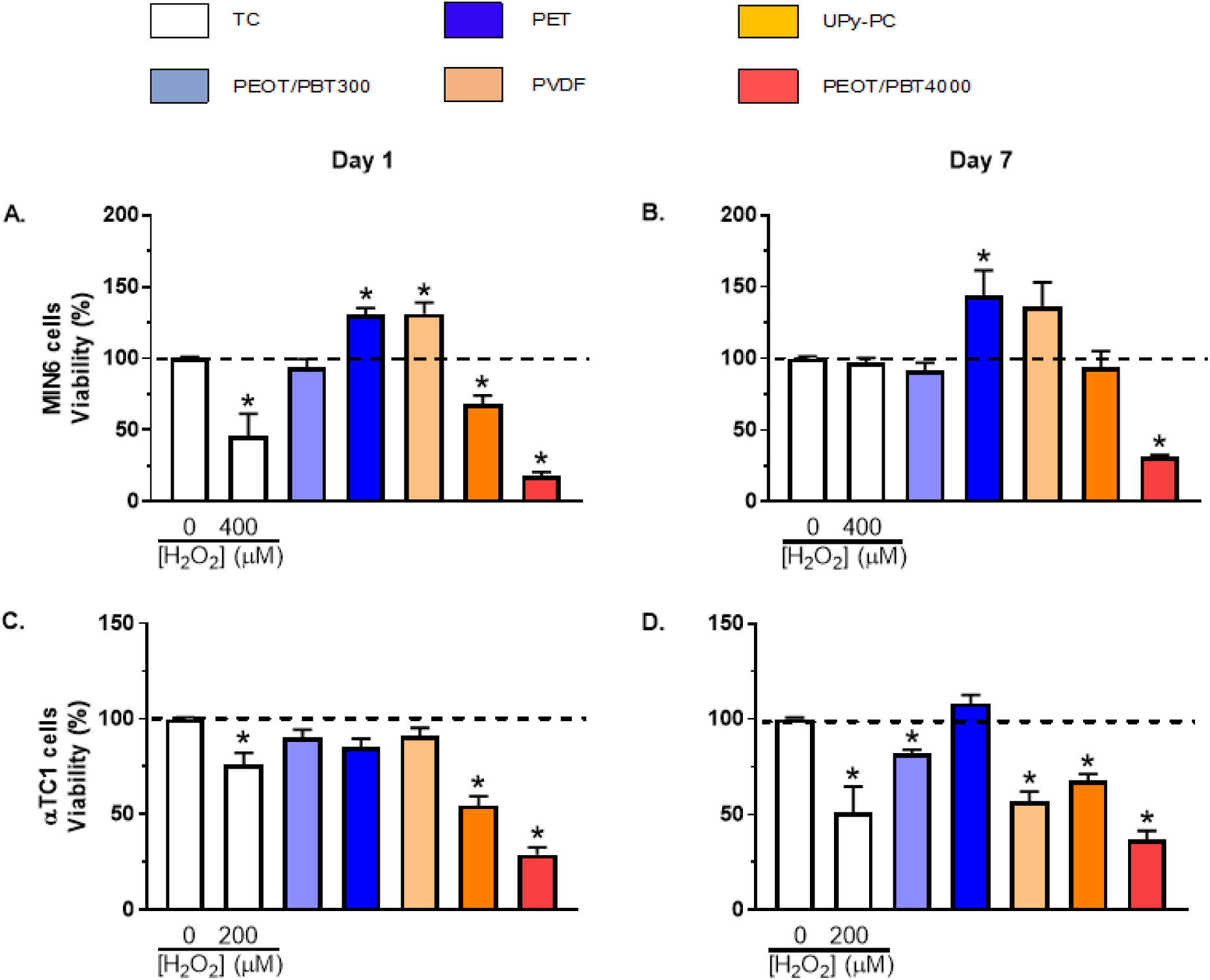
Viability of MIN6 and αTC1 cells cultured on different polymer films. UPy-PC and PEOT/PBT4000 reduced the viability of MIN6 cells on day 1, while PET and PVDF increased viability (A). On day 7, a similar pattern in viability was seen in MIN6 cells exposed to the different biomaterials (B). In αTC1 cells, UPy-PC and PEOT/PBT4000 only decreased the viability on day 1 (C). On day 7, a similar effect was seen and PEOT/PBT300 and PVDF only showed a small decrease in viability (D). Hydrogen peroxide reduced viability, except in MIN6 cells at day 7. N=3 and data are presented as mean ± SEM; * p≤0.05.

**Figure 6:**
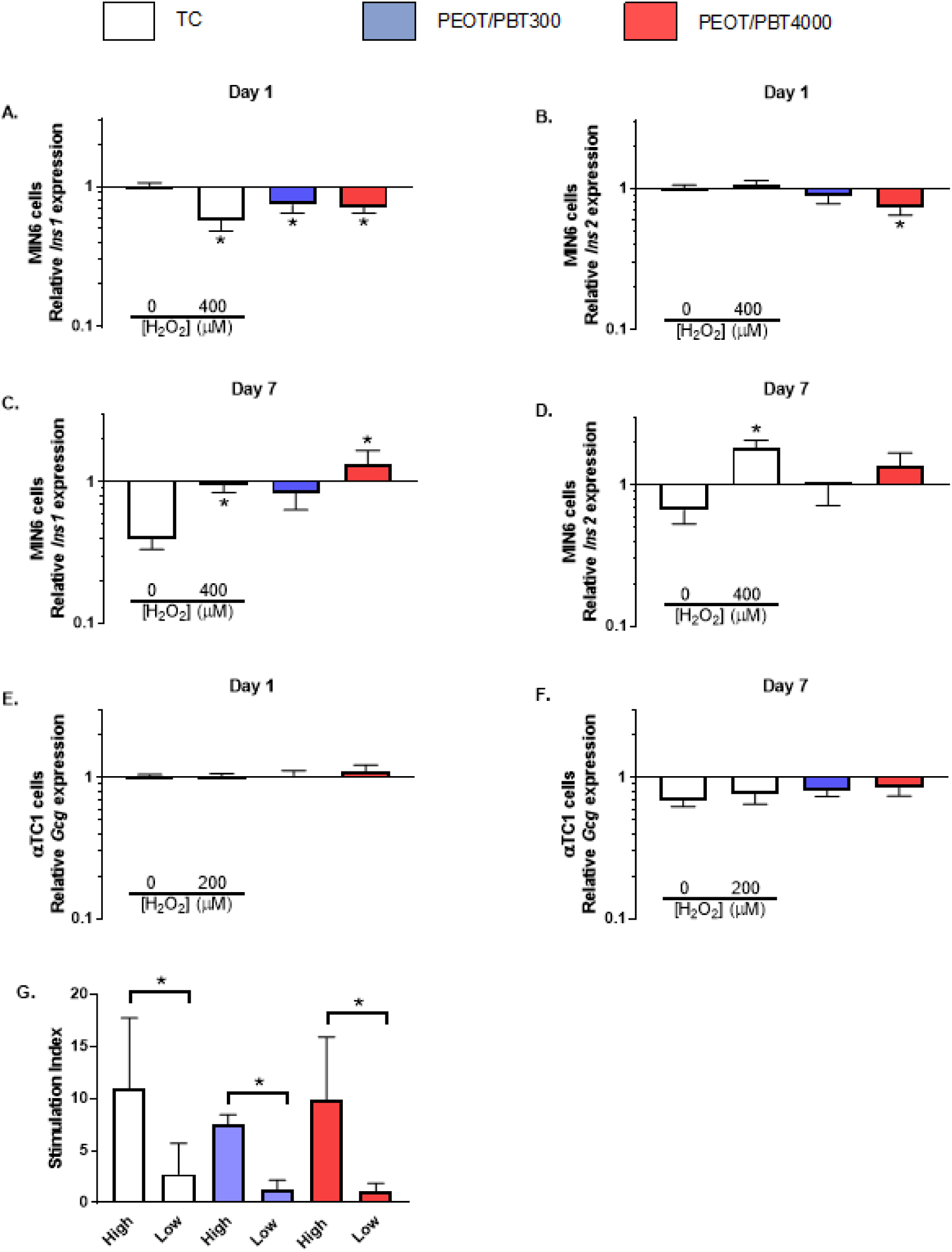
Gene expression levels of insulin-secreting and glucagon-secreting markers in MIN6 and αTC1 cells, respectively, and insulin secretion of human islets cultured on PEOT/PBT300 and PEOT/PBT4000. PEOT/PBT4000 reduced *Ins1* and *Ins2* expression and PEOT/PBT300 *Ins1* expression in MIN6 cells (A–B) on day 1. On day 7 PEOT/PBT4000 increased *Ins1* expression (C–D). *Gcg* expression in αTC1 cells was not significantly different cultured on PEOT/PBT300 and PEOT/PBT4000 compared to TC (E–F) at day 1 and day 7. Hydrogen peroxide (H_2_O_2_) decreased *Ins1* expression, while *Ins2* expression was increased at day 7. *Gcg* expression was not affected by hydrogen peroxide. PEOT/PBT300 and PEOT/PBT4000 did not decrease the stimulation index in human islets (G). N=3 (A–F) and N≥5 (G) and data are presented as mean ± SEM; * p≤0.05.

### PEOT/PBT300 did not influence insulin secretion

To determine if the level of oxidative stress induced by the biomaterials affected the insulin-secreting function of beta cells and glucagon-secreting function of alpha cells, we quantified mRNA levels of several important marker genes: *Ins1* and *Ins2* in MIN6 cells, and *Gcg* in αTC1 cells cultured on PEOT/PBT300 and PEOT/PBT4000. We selected these two polymers because they consistently induced low and high levels of oxidative stress, respectively (Figure 3 and 4). *Ins1* and *Ins2* transcript levels were significantly decreased (approximately 0.76 compared to the TC control of 1.00) at day 1 in MIN6 cultured on both PEOT/PBT300 and PEOT/PBT4000 (Figure 6A–B). At day 7, *Ins1* and *Ins2* expression was not affected on PEOT/PBT300 (Figure 6C), while PEOT/PBT4000 increased *Ins1* expression to 1.36 (Figure 6D). *Gcg* expression in αTC1 cells at day 1 was not changed by culturing them on PEOT/PBT300 and PEOT/PBT4000 compared to TC (Figure 6E). At day 7, both PEOT/PBT300 and PEOT/PBT4000 decreased *Gcg* expression to 0.81 and 0.86 relative to TC at day 1 (Figure 6E–F). Culturing αTC1 on TC also had an effect at day 7, where *Gcg* expression was decreased to 0.70 (Figure 6G).

With these results in the two cell lines, we were prompted to determine whether PEOT/PBT300 and PEOT/PBT4000 would affect the insulin secretion of primary human islets, a critical function for any biomaterial used in an encapsulation device. In contrast to the *Ins1* and *Ins2* transcript downregulation in MIN6 cells, insulin secretion of primary human islets cultured on these two biomaterials was similar to that of islets cultured on control TC (Figure 6G). These results indicate that oxidative stress levels conferred by these polymers did not affect normal insulin secretion but decreased insulin transcription (Figure 6A–F), which may have long-term effects that are not observable in the current studies.

## Discussion

In this study, biomaterial properties, angiogenesis-related gene expression, oxidative stress levels, expression of endogenous antioxidant, beta and alpha cell viability and functionality, and insulin secretion are considered to select a biomaterial with suitable properties for use in an islet encapsulation device.

Multiple biomaterial properties can influence the mechanical characteristics of biomaterials, including their chemical structure, thickness, (pore) size, and the manufacturing parameters. Taking into account some variation related to differences in manufacturing, we could confirm that the mechanical characteristics of all biomaterials we measured were in agreement with results from previous studies [42-45]. These characteristics could mainly be attributed to the chemical structure, where the block copolymers comprising soft, hydrophilic PEO segments are crosslinked with hard, semicrystalline PBT segments providing both strength and elasticity compared to a polymer made from only one monomer subunit like PVDF [45]. In addition, the hydrophobicity of PVDF could be explained by its fluorine group [46]. We found that PEOT/PBT300 and PEOT/PBT4000 were both elastic and resistant to breakage, making them suitable for implantation and immunoprotection. Knowing that scaffold stiffness has been shown to influence cell behavior by modulating the extracellular matrix and changing the islet niche [14-16], we measured the Young’s moduli of the materials. PEOT/PBT300 and UPy-PC had a modulus of 482 KPa and 234 KPa, respectively, which is close to the Young’s modulus of the liver (the main implantation site for CIT [47]) of 10.5 KPa and pancreas of 1.4-4.4 KPa [48, 49]. In addition, PEOT/PBT300 and PEOT/PBT4000 are more hydrophilic compared to PVDF and less hydrophilic than PET, a property that will allow the diffusion of hydrophilic molecules like insulin and glucose to the transplanted islets.

In addition to the material properties, we focused on oxidative stress because it decreases islet viability, and because there was a lack of knowledge on the oxidative stress-inducing effect of biomaterials on beta or alpha cell behavior. Hydrogen peroxide (H_2_O_2_) was used as a positive inducer of oxidative stress [50]. We found that αTC1 are more sensitive to H_2_O_2_ than MIN6. In addition, different biomaterials induced different intracellular oxidative stress levels in MIN6 and αTC1 cells. We hypothesize that the underlying mechanism for this is that the biomaterials induce the formation of ions or small molecules of various sizes [51], which in turn induce cellular oxidative stress. Each biomaterial releases substances of different composition and size with a different release profile, which could explain the different effects on the cells. Whether these possibilities or other reasons underlie our observations remain to be elucidated. We expect even more pronounced effects after implantation, because the wound repair process can trigger a calcium flux via gap junctions of neighboring cells, activating the DUOX/lactoperoxidase system to produce H_2_O_2_ meant to kill invaders and attract leukocytes [52, 53]. This immune response will further contribute to the oxidative stress (and inflammation) caused by the biomaterials. Further evidence for the importance of considering oxidative stress comes from a study where a biomaterial was modified with antioxidants, which diminished fibrotic encapsulation upon implantation [54, 55].

Our findings that oxidative stress levels diminished in MIN6 cells over time, whereas they remained high in αTC1 cells (Figure 3), implies that alpha cells have a greater sensitivity to oxidative stress and should thus also be considered in studies that are typically focused on the critical insulin-producing beta cells. This study showed that MIN6 cells alone could adapt to oxidative stress through an increased expression of the endogenous antioxidant heme oxygenase-1, a mechanism that was previously shown to decrease in intracellular oxidative stress [56].

The biomaterial-induced oxidative stress did not decrease insulin secretion from human islets compared to control (Figure 6), which is in agreement with previous findings [57]. However, expression of *Ins1* and *Ins2* in MIN6 cells was significantly decreased by both PEOT/PBT300 and PEOT/PBT4000, while *Gcg* expression in αTC1 cells was not affected, indicating that oxidative stress induced by the biomaterials did affect the transcription of insulin. The measured insulin could also be attributed to short-term compensation by functional beta cells that get subsequently depleted, since not all beta cells are active in a physiological islet [58-60]. Whether reduced transcription of *Ins1* and *Ins2* in MIN6 cells affects insulin secretion at a longer time scale (> 7 days) remains to be determined.

In conclusion, when selecting the most suitable biomaterial for an islet encapsulation device, many characteristics like device design, surgical handling, *in vivo* performance, and the method of fabrication should be taken into account and an optimal biomaterial is going to be a compromise between stress, biomaterial properties, surgical handling, device fabrication and cell behavior. The biomaterial that has the best combination of properties is what we call a pancreatic cell compatible biomaterial. Based on biomaterial properties, angiogenesis-related gene expression, oxidative stress levels, expression of endogenous antioxidant, beta and alpha cell viability and functionality, and insulin secretion, PEOT/PBT300 is a well-qualified candidate for the development of a future islet implantation device. It is elastic and resistant to breakage, making it suitable for implantation and immunoprotection. In addition, it is relatively hydrophilic, enhancing the diffusion of insulin and glucose. Angiogenesis-related genes were not negatively affected, and alpha and beta cell viability was not decreased on this polymer. Importantly, it did not induce oxidative stress or affect insulin secretion. In the future, it would be interesting to investigate PEOT/PBT300 as an implantation device in *in vivo* studies, or to add an antioxidant inducer to an oxidative stress–inducing biomaterial [61, 62] to improve the outcomes of CIT.

## Supplement

**Supplementary Table 1:** Statistical significance of biomaterials properties parallel to the direction of casting (p≤0.05, n.s.: not significant).

| **Peak stress** | PEOT/PBT300 | PET | PVDF | UPy-PC | PEOT/PBT4000 |
| --- | --- | --- | --- | --- | --- |
| PEOT/PBT300 |  | n.s. | * | * | * |
| PET |  |  | * | * | n.s. |
| PVDF |  |  |  | n.s. | * |
| UPy-PC |  |  |  |  | * |
| PEOT/PBT4000 |  |  |  |  |  |
| **Young's modulus** | PEOT/PBT300 | PET | PVDF | UPy-PC | PEOT/PBT4000 |
| PEOT/PBT300 |  | * | * | * | * |
| PET |  |  | * | * | * |
| PVDF |  |  |  | * | * |
| UPy-PC |  |  |  |  | * |
| PEOT/PBT4000 |  |  |  |  |  |
| **Failure stress** | PEOT/PBT300 | PET | PVDF | UPy-PC | PEOT/PBT4000 |
| PEOT/PBT300 |  | * | * | * | * |
| PET |  |  | * | * | * |
| PVDF |  |  |  | n.s. | n.s. |
| UPy-PC |  |  |  |  | n.s. |
| PEOT/PBT4000 |  |  |  |  |  |
| **Failure strain** | PEOT/PBT300 | PET | PVDF | UPy-PC | PEOT/PBT4000 |
| PEOT/PBT300 |  | * | * | * | n.s. |
| PET |  |  | * | * | * |
| PVDF |  |  |  | * | * |
| UPy-PC |  |  |  |  | * |
| PEOT/PBT4000 |  |  |  |  |  |
| **Water contact angle** | PEOT/PBT300 | PET | PVDF | UPy-PC | PEOT/PBT4000 |
| TC | n.s. | * | * | * | n.s. |
| PEOT/PBT300 |  | * | * | * | n.s. |
| PET |  |  | * | * | * |
| PVDF |  |  |  | * | * |
| UPy-PC |  |  |  |  | n.s. |
| PEOT/PBT4000 |  |  |  |  |  |

**Supplementary Table 2:**
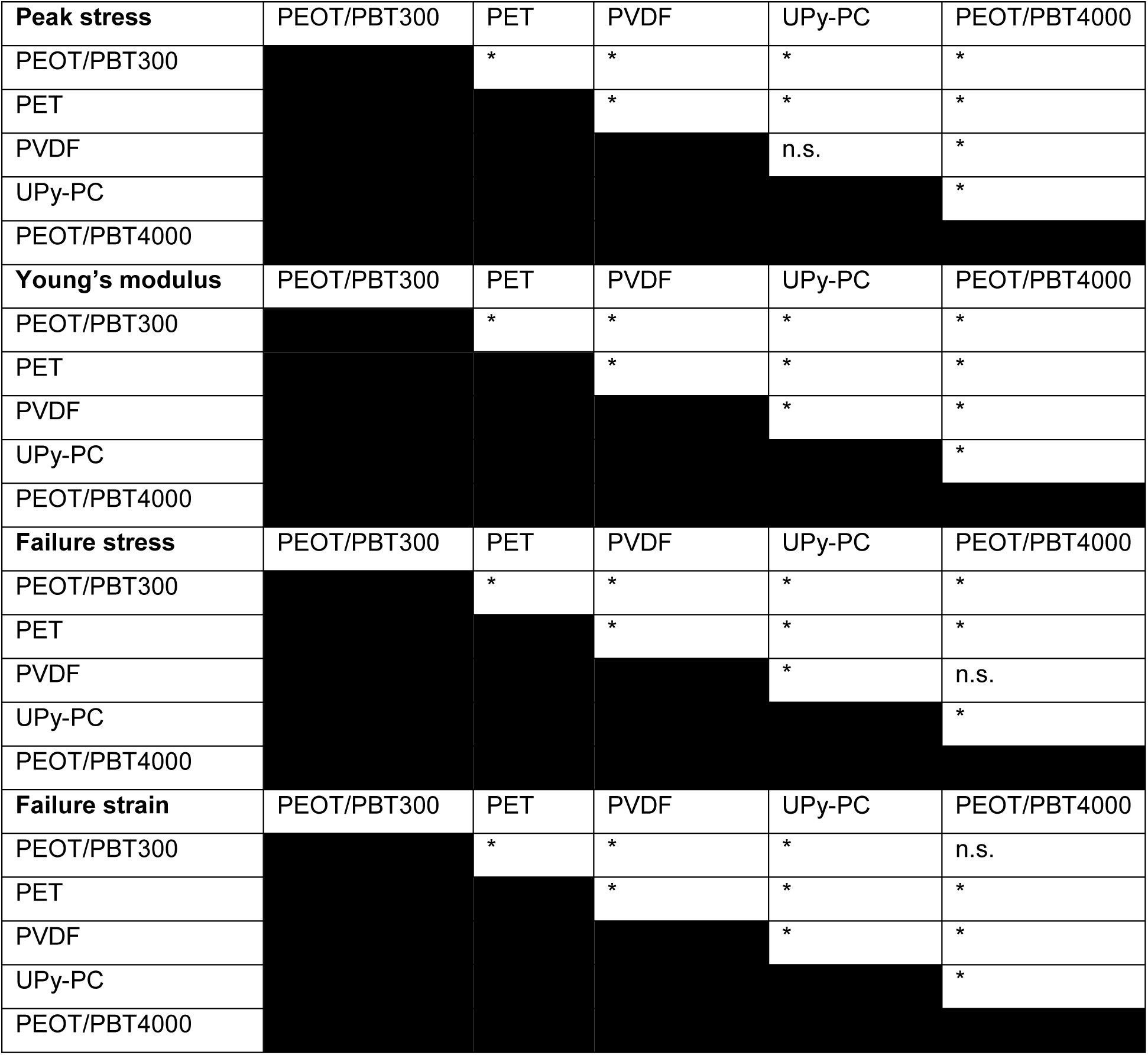
Statistical significance of biomaterials properties perpendicular to the direction of casting (p≤0.05, n.s.: not significant).6

| **Peak stress** | PEOT/PBT300 | PET | PVDF | UPy-PC | PEOT/PBT4000 |
| --- | --- | --- | --- | --- | --- |
| PEOT/PBT300 |  | * | * | * | * |
| PET |  |  | * | * | * |
| PVDF |  |  |  | n.s. | * |
| UPy-PC |  |  |  |  | * |
| PEOT/PBT4000 |  |  |  |  |  |
| **Young's modulus** | PEOT/PBT300 | PET | PVDF | UPy-PC | PEOT/PBT4000 |
| PEOT/PBT300 |  | * | * | * | * |
| PET |  |  | * | * | * |
| PVDF |  |  |  | * | * |
| UPy-PC |  |  |  |  | * |
| PEOT/PBT4000 |  |  |  |  |  |
| **Failure stress** | PEOT/PBT300 | PET | PVDF | UPy-PC | PEOT/PBT4000 |
| PEOT/PBT300 |  | * | * | * | * |
| PET |  |  | * | * | * |
| PVDF |  |  |  | * | n.s. |
| UPy-PC |  |  |  |  | * |
| PEOT/PBT4000 |  |  |  |  |  |
| **Failure strain** | PEOT/PBT300 | PET | PVDF | UPy-PC | PEOT/PBT4000 |
| PEOT/PBT300 |  | * | * | * | n.s. |
| PET |  |  | * | * | * |
| PVDF |  |  |  | * | * |
| UPy-PC |  |  |  |  | * |
| PEOT/PBT4000 |  |  |  |  |  |

**Supplementary figure 1:**
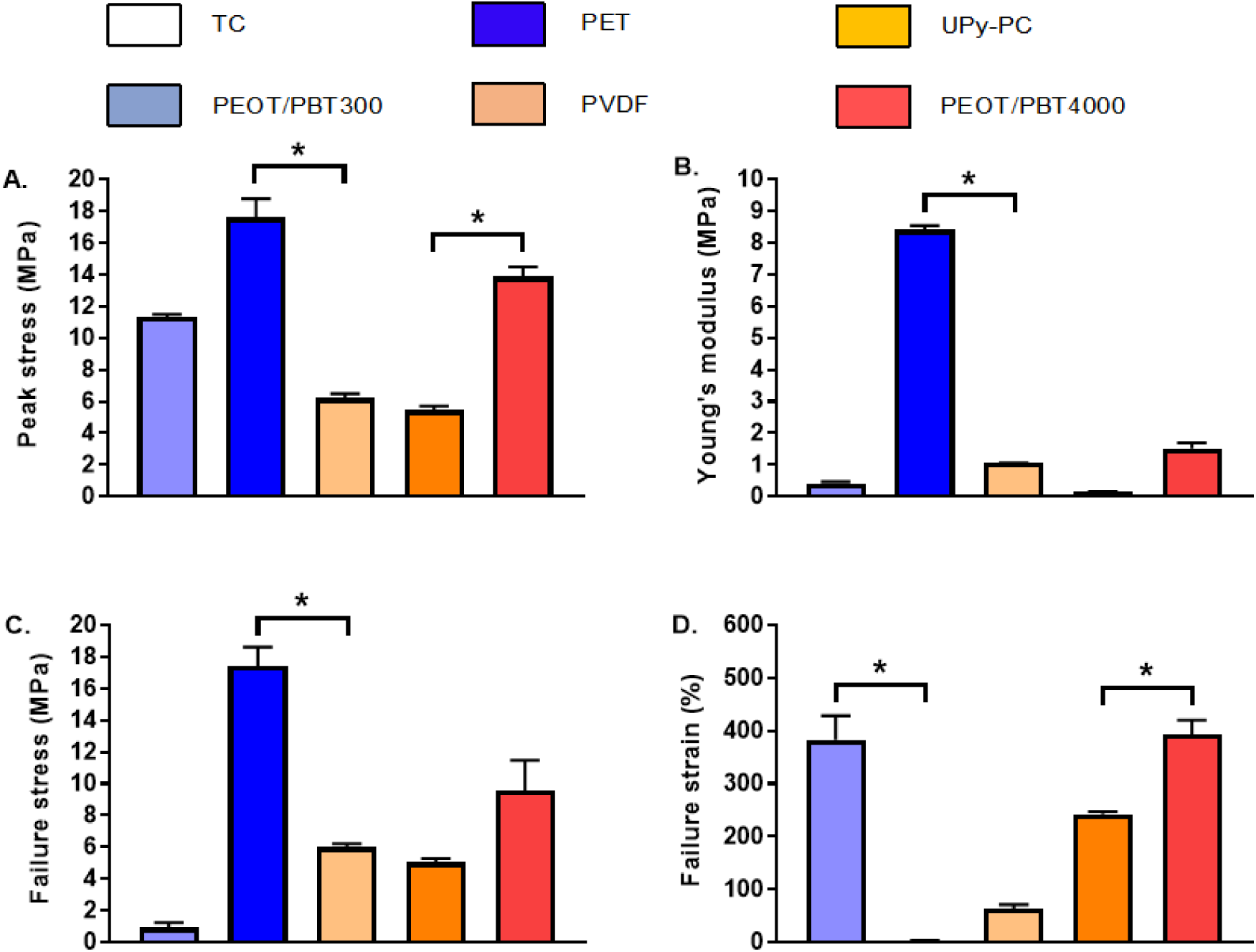
Tensile testing in the direction perpendicular to casting revealed that PET and PEOT/PBT4000 had higher peak stress than the other biomaterials (A). The Young’s modulus and failure stress of PET was higher than the other biomaterials (B–C). PC and PEOT/PBT4000 had the highest failure strain (D). *N*=3 and data are presented as mean ± SEM; * *p* ≤ 0.05.

**Supplementary figure 2:**
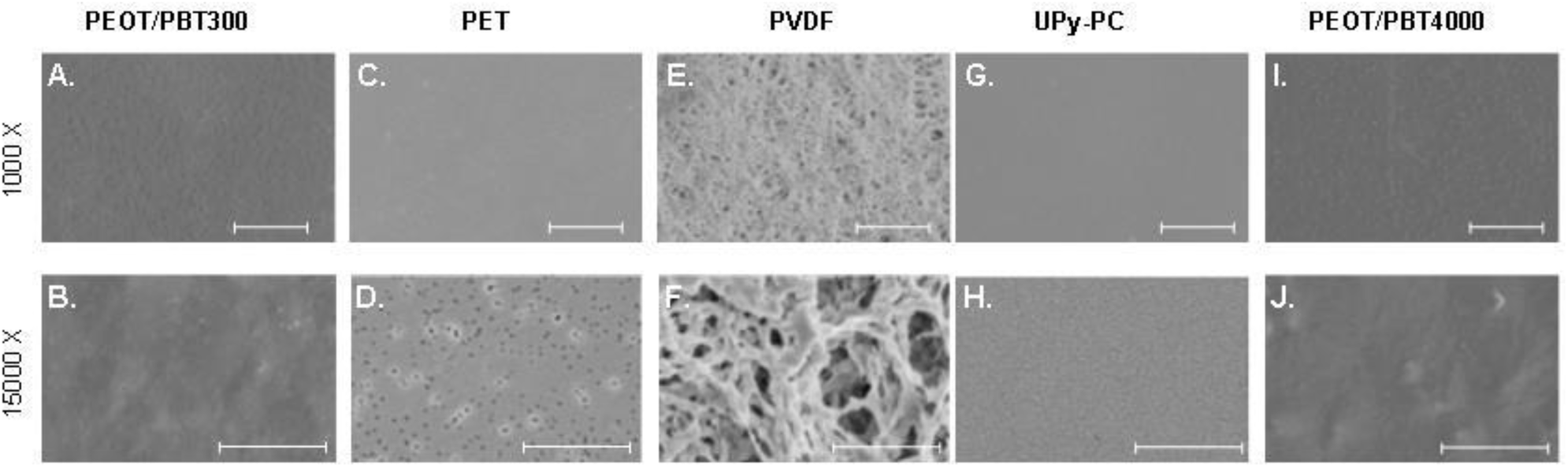
SEM images of PEOT/PBT300, PET, PVDF, UPy-PC, PEOT/PBT4000 on their smooth side showed a different surface structure for the different biomaterials. Bars in the 1000x magnification are 50 μm and bars in the 15000x magnification are 5 μm.

**Supplementary figure 3:**
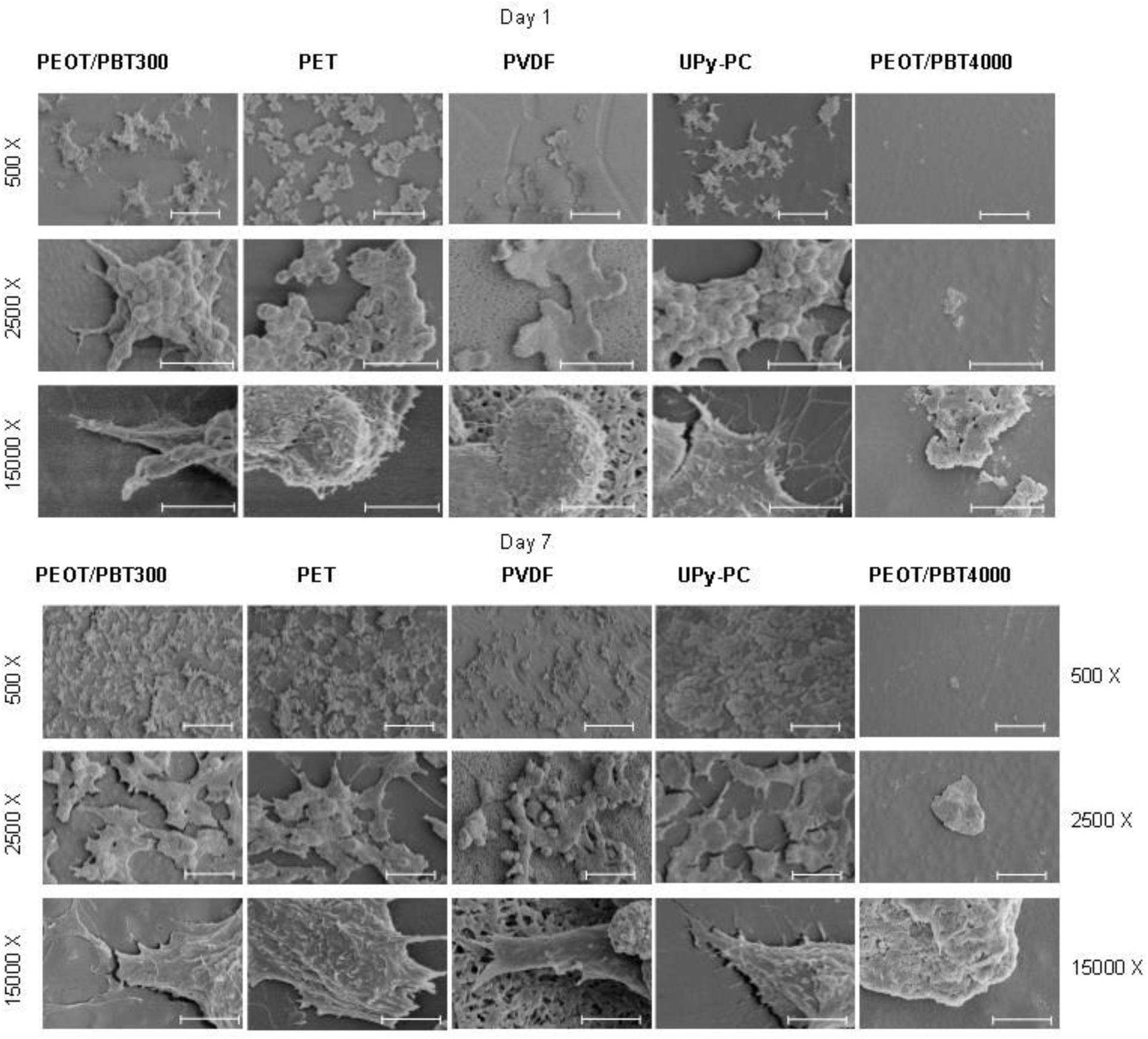
SEM images of MIN6 cells on PEOT/PBT300, PET, PVDF, UPy-PC, PEOT/PBT4000 for 7 days showed no differences in cell morphology. PEOT/PBT4000 had fewer adherent cells. Bars in the 1000x, 2500x and 15000x magnification are 100, 30 and 5 μm respectively.

**Supplementary figure 4:**
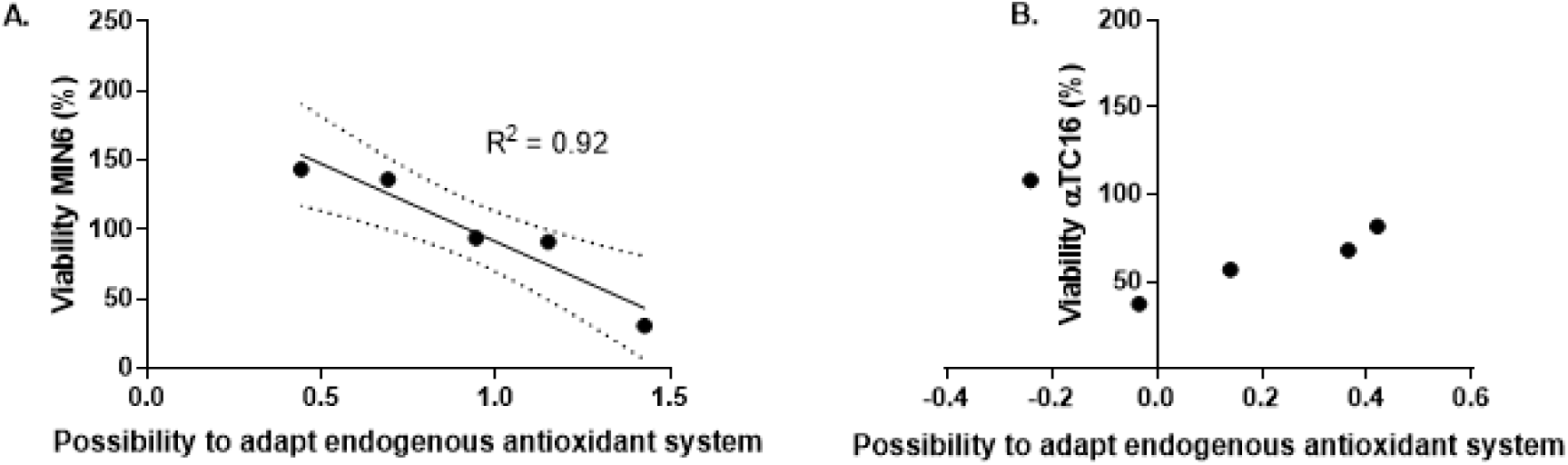
The higher the ability to induce *Hmox1* expression over time from day 1-7 on the different biomaterials, the higher the viability of MIN6 cells on that specific biomaterial (A), while in αTC1 no correlation between *Hmox1* expression over time from day 1-7 on the different biomaterials and the viability was found (B).

## Acknowledgements

This project/research has received funding from the European Research Council (ERC) under the European Union’s Horizon 2020 research and innovation programme (grant agreement No 694801). In addition, we would like to thank Hang Nguyen (MERLN/M4I, Maastricht University, the Netherlands) for her help revising the manuscript.

